# A dynamic interaction between CD19 and the tetraspanin CD81 controls B cell co-receptor trafficking

**DOI:** 10.1101/768275

**Authors:** Katherine J. Susa, Tom C. M. Seegar, Stephen C. Blacklow, Andrew C. Kruse

**Affiliations:** Department of Biological Chemistry and Molecular Pharmacology, Blavatnik Institute, Harvard Medical School, Boston MA 02115; Dana Farber Cancer Institute, Department of Cancer Biology, Boston MA 02215

## Abstract

CD81 and its binding partner CD19 are core subunits of the B cell co-receptor complex. While CD19 is a single-pass transmembrane protein belonging to the extensively studied Ig superfamily, CD81 belongs to a conserved but poorly understood family of four-pass transmembrane proteins called tetraspanins. These functionally diverse proteins play important roles in a wide variety of different organ systems by controlling protein trafficking and other cellular processes. Here, we show that CD81 relies on its ectodomain to control trafficking of CD19 to the cell surface. Moreover, the anti-CD81 antibody 5A6, which binds selectively to activated B cells, recognizes a conformational epitope on CD81 that is masked when CD81 is in complex with CD19. Mutations of CD81 in this contact interface suppress its CD19 surface-export activity. Taken together, these data indicate that the CD81 - CD19 interaction is dynamically regulated upon B cell activation, suggesting that this dynamism can be exploited to regulate B cell function. These results are not only important for understanding B cell biology, but also have important implications for understanding tetraspanin function more generally.

## INTRODUCTION

The tetraspanins constitute a 33-member family of transmembrane proteins in humans. Although poorly understood, tetraspanins play a critical role in mammalian physiology, functioning in nearly all cell types and regulating distinct processes such as control of cell morphology, cell adhesion, protein trafficking, and signal transduction (Hemler, 2008). Tetraspanins are thought to achieve their biological functions through interactions with partner proteins, leading to formation of signaling complexes and modulation of signaling activity (Hemler, 2005). Members of the tetraspanin protein family share an overall domain organization consisting of four transmembrane segments, a small extracellular loop (SEL), a large extracellular loop (LEL) containing a conserved Cys-Cys-Gly (CCG) motif, a short cytoplasmic N-terminal region, and a C-terminal cytoplasmic tail. However, the molecular details of how these domains mediate complex formation with partner proteins to regulate their trafficking and signaling remain unclear.

CD81, the first tetraspanin identified, was discovered as the target of an antiproliferative antibody called “5A6”, which inhibits the growth of B cell lymphoma cell lines (Oren et al., 1990). CD81 plays a critical role in regulating B cell receptor (BCR) signaling as one subunit of the B cell co-receptor complex, which also includes CD19 and CD21 (Carter and Barrington, 2004). Within this complex, CD81 directly interacts with CD19, a single-pass transmembrane protein that establishes the threshold for both BCR dependent and independent signaling. Stimulation of CD19 lowers the signaling threshold needed for both antigen-independent and antigen-dependent activation of B cells by several orders of magnitude, and this signaling is critical for the function of the humoral immune response (Carter and Fearon, 1992; Gauld et al., 2002). Not surprisingly, aberrant CD19 signaling is implicated the development of B cell malignancies, autoimmunity, and immunodeficiency (Barrena et al., 2005; Mei et al., 2012; Yazawa et al., 2005; van Zelm et al., 2006). CD19 is also the target of chimeric antigen receptor expressing T cells now used clinically in the treatment of B cell malignancies (Brentjens et al., 2013; Grupp et al., 2013; Kalos et al., 2011; Kochenderfer et al., 2012; Porter et al., 2011).

Despite the importance and therapeutic relevance of the B cell co-receptor, surprisingly little is known about how CD81 engages CD19 to regulate its trafficking or signaling activity. The association between CD19 and CD81 was first detected using co-immunoprecipiation studies (Bradbury et al., 1992). Later, genetic evidence revealed that defects in complex formation between CD19 and CD81 result in severe deficiencies in B cell function. For example, three independent lines of CD81-null mice showed reduced CD19 surface expression accompanied by defects in B cell function such as weaker early antibody responses, impaired B cell proliferation, and reduced calcium influx following B cell activation (Maecker and Levy, 1997; Miyazaki et al., 1997; Tsitsikov et al., 1997). Additionally, there are human cases of common variable immune deficiency (CVID) in which CD19 expression on B cells is suppressed by homozygous truncations in the CD81 gene (van Zelm et al., 2010).

It has been proposed that CD81 has two key roles as a B cell co-receptor subunit. First, it is thought to chaperone CD19 through the secretory pathway to the plasma membrane (Braig et al., 2016; Shoham et al., 2003). Second, it may also serve as a regulator of B cell signaling by controlling the localization of CD19 at the plasma membrane during B cell activation (Mattila et al., 2013). The molecular mechanisms by which CD81 carries out both trafficking of CD19 and regulation of its signaling activity, however, remain unclear.

Here, we find that CD81 uses its ectodomain to bind CD19 and to promote the export of CD19 to the cell surface. Remarkably, the anti-CD81 antibody 5A6, which binds selectively to activated B cells, recognizes an unusual conformational epitope on CD81 that is masked when CD81 is in complex with CD19, but which becomes accessible upon B cell activation. These findings suggest that the CD81 - CD19 interaction is dynamic and is linked to the B cell activation state.

## RESULTS

### The large extracellular loop of CD81 is required to promote CD19 export to the cell surface

Prior studies have suggested that the first TM helix of CD81 is the main specificity determinant for trafficking of CD19 to the cell surface (Shoham et al., 2006), but this claim has also been called into question (Berditchevski and Odintsova, 2007). To determine which regions of CD81 are necessary for trafficking CD19 to the cell surface, we established a HEK293T cell line in which CD81 was knocked out with CRISPR-Cas9 and used flow-cytometry to test the ability of CD81 or CD81 chimeric proteins to enhance delivery of CD19 to the cell surface. We created chimeras of CD81 with CD9 (the tetraspanin most similar to CD81 in sequence) and with Tspan15 (a divergent tetraspanin from *C. elegans*). The chimeras have domain swaps of the small extracellular loop, large extracellular loop, or first transmembrane helix **(Figure 1A)**. Expression of CD81 chimeras was confirmed by flow cytometry **(Supplementary Figure 1)**.

**Figure 1:**
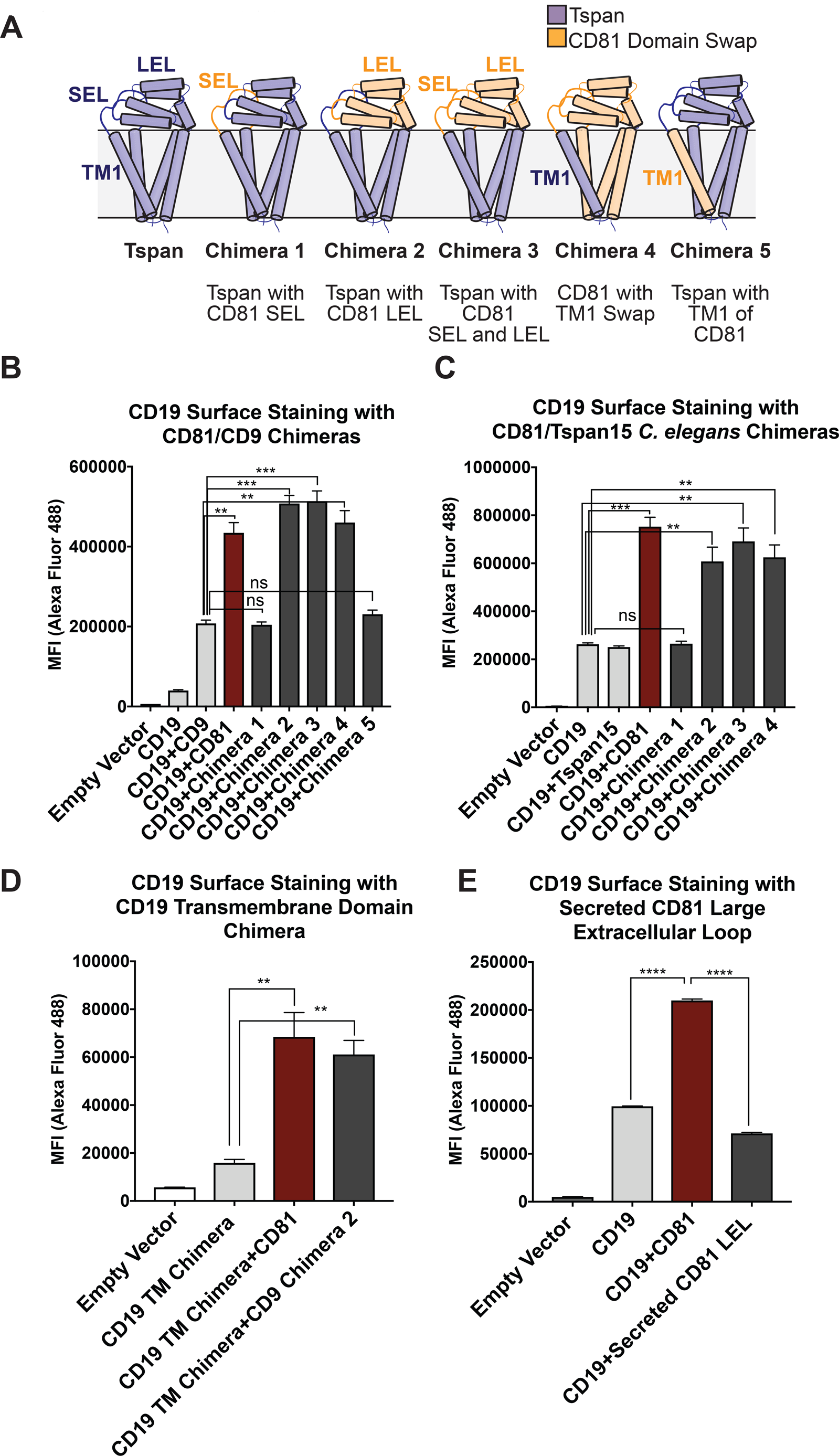
CD81 chimera design and CD19 Export Assay. **(A)** Design of CD81 Chimeras used in export assay experiments. **(B)** Export assay with CD81/CD9 chimeras. **(C)** Export assay with CD81/Tspan15 *C. elegans* chimeras. **(D)** Export assay with CD19/ σ1 receptor transmembrane domain chimera. **(E)** Export assay with a secreted construct of the CD81 large extracellular loop. For the data in panels **B** - **E**, surface CD19 was detected by flow cytometry using an Alexa 488-coupled anti-CD19 antibody. Each figure represents three independent experiments. Error bars represent mean ± SEM. Statistical analysis was performed in GraphPad Prism using an unpaired two-tailed t test. **p < 0.01; ***p < 0.001, ****p<0.0001.

Whereas cells transfected with CD19 alone only show a small amount of surface staining **(Figure 1B, 1C)**, co-transfection of CD19 with wild-type CD81 results in a two to four-fold enhancement in CD19 surface staining. Chimeras that retain the large extracellular loop of CD81 stimulate the same increase in CD19 surface staining as wild type CD81, but chimeras lacking the large extracellular loop of CD81 do not, indicating that the large extracellular loop is necessary for CD19 surface export. Replacement of the first TM helix of CD81 with that of CD9 or Tspan15 also supports the same increase in surface staining as wild type CD81, whereas replacement of the first TM helix of CD9 with that of CD81 fails to increase surface staining of CD19, revealing that the first TM helix of CD81 in the context of a CD9 backbone is not sufficient to support the trafficking of CD19 **(Figure 1B, 1C)**.

To assess whether surface export of CD19 depends upon the presence of its native TM region, we created a CD19 chimera in which its native TM was replaced with that of the σ1 receptor (Schmidt et al., 2016a). CD81 exports the CD19/σ1 chimera to the cell surface as effectively as it exports wild-type CD19, indicating that it is association of the ectodomains that promotes the trafficking of CD19 to the cell surface **(Figure 1D)**. A secreted form of the CD81 large extracellular loop, however, does not promote increased CD19 surface expression, suggesting that membrane tethering plays an important role in CD19 surface delivery by increasing the effective concentrations of the two proteins for each other **(Figure 1E)**.

### The epitope of the 5A6 CD81 antibody is masked when CD81 is in complex with CD19

To further characterize the CD19-CD81 complex, we constructed a fusion protein in which the C-terminus of CD19 is directly connected to the N-terminus of CD81 with a short intervening linker **(Figure 2A)**. To assess the integrity of this fusion protein, we evaluated its abundance on the cell surface by flow cytometry, and examined the reactivity of the fusion protein with a panel of anti-CD19 and anti-CD81 antibodies (Nelson et al., 2018). The abundance of the CD19-CD81 fusion protein on the cell surface is comparable to that observed when full-length CD19 is co-expressed with wild type CD81, indicating that CD81 in the fusion protein is functional in trafficking CD19 to the cell surface **(Figure 2B)**. Moreover, four different anti-CD19 antibodies recognize the fusion protein **(Supplementary Figure 2)**, as do three anti-CD81 antibodies, providing further evidence that both CD19 and CD81 are properly folded in the context of the fusion protein. Surprisingly, however, one anti-CD81 antibody, called 5A6 (Levy et al., 2017; Oren et al., 1990), showed significantly decreased binding of the fusion protein compared to the other anti-CD81 antibodies **(Figure 2C)**. Although all four antibodies bind the large extracellular loop of CD81, only 5A6 is unable to detect the CD19-CD81 fusion protein, suggesting that its epitope overlaps with the region(s) of CD81 that contact CD19 in the native complex (Nelson et al., 2018). A prior co-immunoprecipitation experiment also showed that the 5A6 antibody cannot be used to pull down the components of CD21/CD19 complex in a B cell line, providing further evidence the 5A6 epitope is masked by CD19 (Matsumoto et al., 1993).

**Figure 2:**
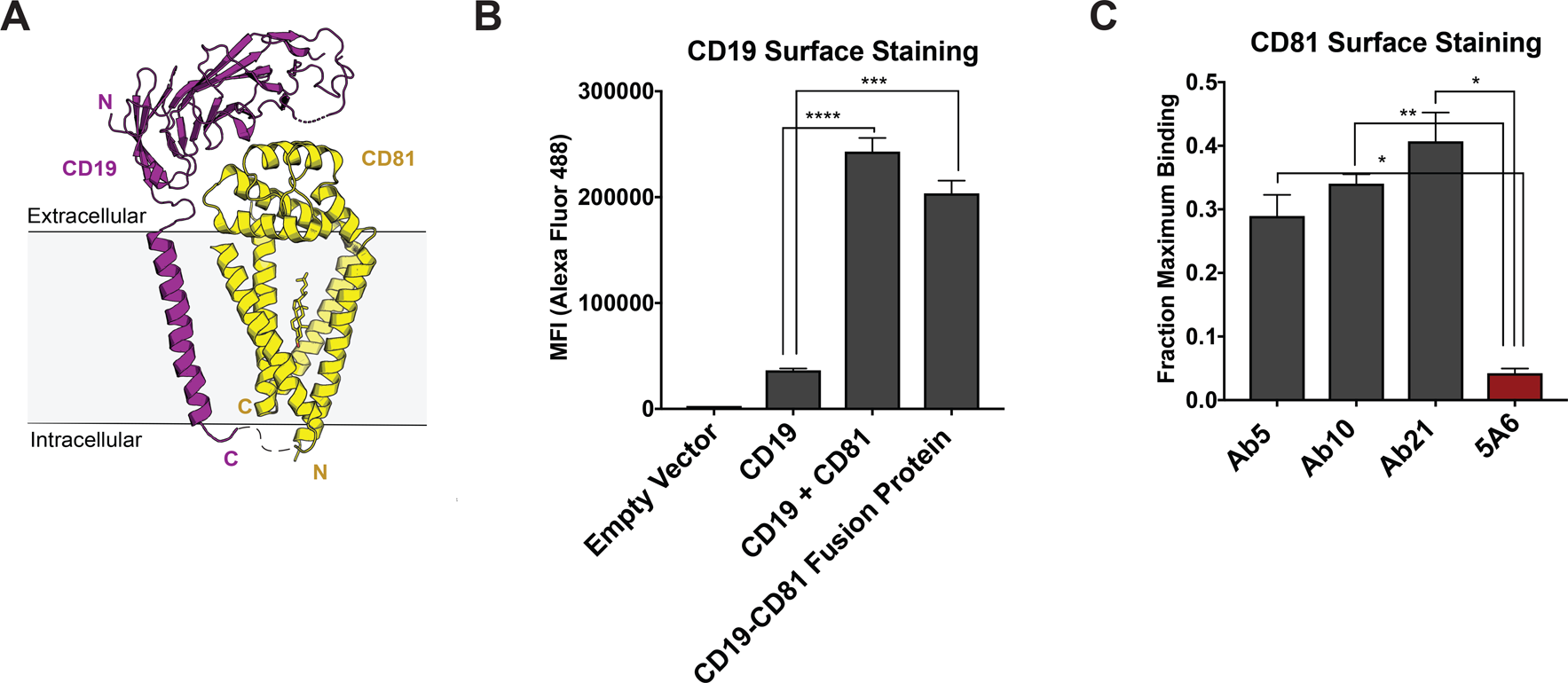
Design and evaluation of a CD19-CD81 fusion protein. **(A)** Cartoon representing the designed CD19-CD81 fusion protein. The model was created based on known structures of the CD19 ectodomain (PDB 6AL5) and CD81 (PDB 5TCX). A short Gly-Ser linker (dashed lines) connects P329 of the intracellular portion of CD19 to the N-terminus of CD81. **(B)** Analysis of CD19 surface staining in CD81-null cells expressing the CD19-CD81 fusion protein. Surface staining for the CD19-CD81 fusion is compared to staining of cells expressing only CD19, and to staining of cells expressing both CD19 and CD81, using an Alexa 488-coupled anti-CD19 antibody. **(C)** Binding of various CD81 antibodies to the CD19-CD81 complex, analyzed by flow cytometry. “Fraction maximum binding” was calculated by dividing the average MFI of antibody bound to CD19-CD81 by the average MFI of antibody bound to CD81. An anti-human IgG-Alexa 488 secondary antibody was used to detect CD81 antibody bound to the cell surface. For the data in panel B and C, each figure represents three independent experiments and error bars represent mean ± SEM. Statistical analysis was performed in GraphPad Prism using an unpaired t test. **p < 0.01; ***p < 0.001, ****p<0.0001.

### 5A6 Binds to helices C and D of CD81

To gain insight into the molecular basis underlying the unique reactivity of the 5A6 antibody, we determined the structure of the 5A6 F_ab_ in complex with the large extracellular loop (LEL) of CD81 to 2.4 Å resolution using x-ray crystallography (Table 1). The overall architecture of the CD81 LEL has five helices, with the A, B and E helices as a stalk and helices C and D capping the “top” face. The 5A6 F_ab_ binds CD81 at an epitope derived almost exclusively from helices C and D **(Figure 3A, B)**, burying a total of 1522 A^2^ of solvent accessible surface area. The paratope of 5A6 is derived from all three heavy chain complementarity-determining regions (CDRs; residues 31-35, 50-66, 99-108) and from the first two light light-chain CDRs (residues 24-40 and 55-61).

**Table 1:** X-ray crystallography data collection and refinement statistics. Refined coordinates and structure factors are deposited in the Protein Data Bank under accession code 6U9S.

| **Data Collection** | **5A6-CD81 LEL** |
| --- | --- |
| Wavelength (Å) | 0.9792 |
| Space Group | P 2<sub>1</sub> 2<sub>1</sub> 2<sub>1</sub> |
| Number of crystals | 1 |
| Unit cell dimensions |  |
| *a, b, c* | 40.003, 96.858, 297.091 |
| $\alpha, \beta, \gamma$ (°) | 90, 90, 90 |
| Resolution (Å) (last shell) | 49.5-2.4 (2.54-2.4) |
| No. of reflections (total/unique) | 293920/46497 |
| Completeness (%) (last shell) | 99.1 (96.8) |
| $\langle I/\sigma(I) \rangle$ (last shell) | 7.94 (0.46) |
| R<sub>meas</sub> (%) (last shell) | 20.5% (355.7%) |
| CC<sub>1/2</sub> (%) (last shell) | 99.5 (16.6) |
| Multiplicity | 6.3 |
| **Refinement** |  |
| Number of atoms (protein/solvent) | 7974/358 |
| R<sub>work</sub>/R<sub>free</sub> (%) | 21.44/28.08 |
| R.M.S. deviation (Å) |  |
| Bond length | 0.003 |
| Bond angles | 0.537 |
| Ramachandran statistics |  |
| Favored | 96.94 |
| Allowed | 3.06 |
| Outliers | 0.00 |

**Figure 3:**
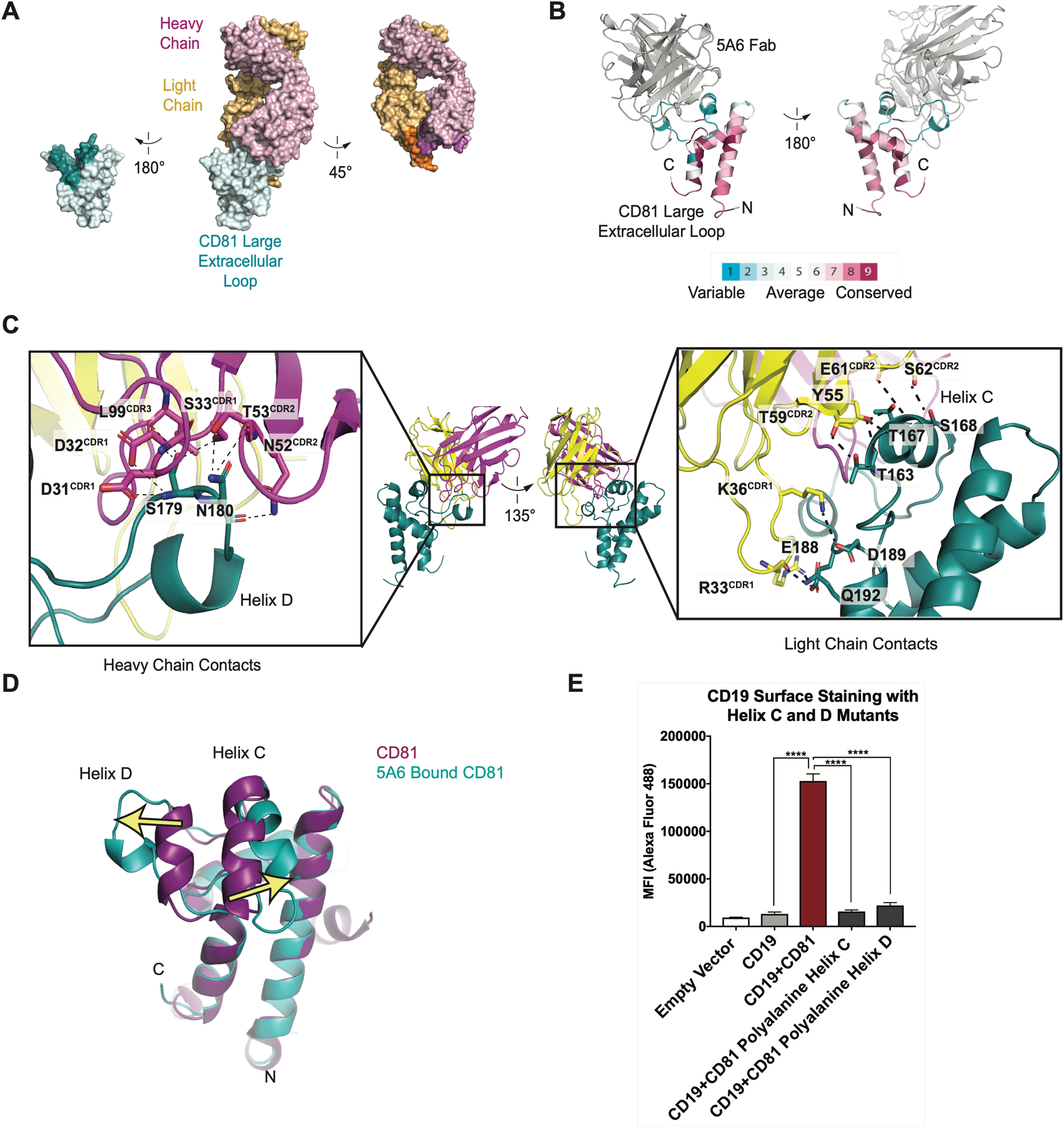
Structure of the 5A6 F_ab_-CD81 Large Extracellular Loop Complex (PDB 6U9S). **(A)** Surface representation of the 5A6-CD81 complex. CD81 is blue, the 5A6 F_ab_ light chain is yellow, and the heavy chain is magenta. Residues at the binding interface are colored in a darker shade. **(B)** CD81 colored by evolutionary conservation score. **(C)** 5A6 F_ab_-CD81 binding interface. Heavy chain (left panel) and light chain (right panel) contacts are shown. Hydrogen bonding interactions are indicated with dotted lines. **(D)** Structural superposition of 5A6 bound CD81 on full length CD81. Arrows indicate positional shifts of the variable helices C and D in the 5A6-bound structure. **(E)** CD19 Export Assay with Helix C and D Mutants. Surface CD19 was detected by flow cytometry using an Alexa 488-coupled anti-CD19 antibody. Expression of helix C and D mutants was confirmed by flow cytometry (**Supplementary Figure 1**). For the data in panel E, error bars represent mean ± SEM of three independent experiments. Statistical analysis was performed in GraphPad Prism using an unpaired t test. **p < 0.01; ***p < 0.001, ****p<0.0001.

The most striking feature of the contact interface is the large-scale rearrangement of the CD81 C and D helices, which splay apart in the structure of the complex (**Figure 3D**). Helix C moves outward by approximately 7 Å, and Helix D unravels almost completely and moves outward by approximately 11 Å, compared to its position in the structure of free full-length CD81. This structural rearrangement occurs because the heavy chain of 5A6 inserts its CDR3 loop between the helices, allowing it to form polar contacts with S179 and N180. Additional key interactions at the F_ab_-CD81 interface include extensive light-chain contacts with helix C of CD81 (**Figure 3C**). Among these interactions are hydrogen-bonds between the side chain hydroxyl group of F_ab_ residue Y55 with the T167 side chain hydroxyl, the T167 backbone amide, and the T163 backbone carbonyl of CD81. Side chain hydrogen bonding interactions are also present between T59 of the F_ab_ and T163 of CD81, and between S62 of the F_ab_ and S168 of CD81. The light chain of 5A6 also contacts three residues at the start of Helix E, forming a hydrogen bonding network with residues E188, D189, and Q192 of CD81.

### Helix C and D of CD81 Mediate CD19 Complex Formation

The binding of 5A6 to CD81 results in substantial conformational changes of Helices C and D in CD81. These two helices form a solvent-exposed, low polarity region in the CD81 large extracellular loop, and evolutionary analysis reveals that this region is highly variable among the different proteins of the tetraspanin family **(Figure 3B)**. Both molecular dynamics simulations and NMR studies suggest the Helix D of CD81 is the most flexible region of the large extracellular loop (Rajesh et al., 2012; Schmidt et al., 2016b). In molecular dynamics simulations, Helices A, B, and E from the large extracellular loop of CD81 retain their alpha helical structure, but Helices C and D are more labile and show a tendency to lose alpha-helicity (Schmidt et al., 2016b).

Because the 5A6 antibody is non-reactive with the CD19-CD81 fusion protein, and because its epitope consists primarily of the CD81 C and D helices, we hypothesized that the C/D helix region of CD81 is essential for CD19 binding. To test this idea, we introduced alanine substitutions in helix C or D and measured the effect of these mutations on export of CD19 to the cell surface using our flow cytometry assay. Mutation of either helix C or D to polyalanine results in decreased trafficking of CD19 to the cell surface, strongly suggesting that each helix contributes to CD19-CD81 complex formation **(Figure 3E)**.

### The CD19-CD81 complex dissociates in activated B cells

Although CD81 has a clear role in trafficking CD19 to the cell surface, its function at the B cell membrane remains unclear. Tetraspanins are thought to organize receptors and associated signaling proteins in functional microdomains in the plasma membrane, thereby regulating receptor signaling and their associated signaling pathways. Super-resolution microscopy suggests that CD19 may be compartmentalized in the B cell membrane by CD81 to regulate signaling through the BCR, but the mechanistic details of how CD81 regulates the localization of CD19 remain poorly understood (Mattila et al., 2013; Zuidscherwoude et al., 2015).

To address this question, we used antibodies with different CD81 binding epitopes to probe the dynamics of the CD19-CD81 complex on primary human B cells in response to B cell activation. We isolated primary human B cells from a fresh leuko-reduction collar and activated them with an anti-B cell receptor (BCR) antibody. Anti-BCR antibody treatment resulted in increased CD69 and CD86 at the cell surface when compared with resting cells, confirming activation of the antibody-stimulated cells **(Figure 4A)**. We then compared the surface staining of CD19 and CD81 in the resting and activated states, using anti-CD81 antibodies with different epitopes to distinguish free CD81 from CD19-associated CD81. CD19 showed no difference in surface staining between resting and activated cells **(Figure 4B)**. When surface CD81 is detected using Ab21 (Nelson et al., 2018), which recognizes both free and complexed CD81 (**Supplementary Figure 3**), there is no difference between resting and activated B cells **(Figure 4B, 4C)**, but when CD81 is detected using the 5A6 antibody, which selectively recognizes free CD81, there is a two-fold increase in surface staining on activated B cells **(Figure 4B, 4C)**. Western blotting of whole cell lysates from resting and activated cells with Ab21 and 5A6 revealed no significant difference in protein levels, indicating that the increase in CD81 surface staining is not due to increased production of CD81 in activated B cells **(Figure 4D)**but instead reflects a change in the accessibility of the 5A6 epitope. This finding indicates that CD81 either undergoes a conformational change or dissociates from CD19 to expose the epitope upon B cell activation. To distinguish between these two possibilities, we immunopurified CD19 from resting and activated primary B cells. Immunoprecipitation of CD19 recovered much more CD81 from resting B cells than it did from activated B cells, indicating that the CD19-CD81 complex dissociates in activated B cells **(Figure 4E)**.

**Figure 4:**
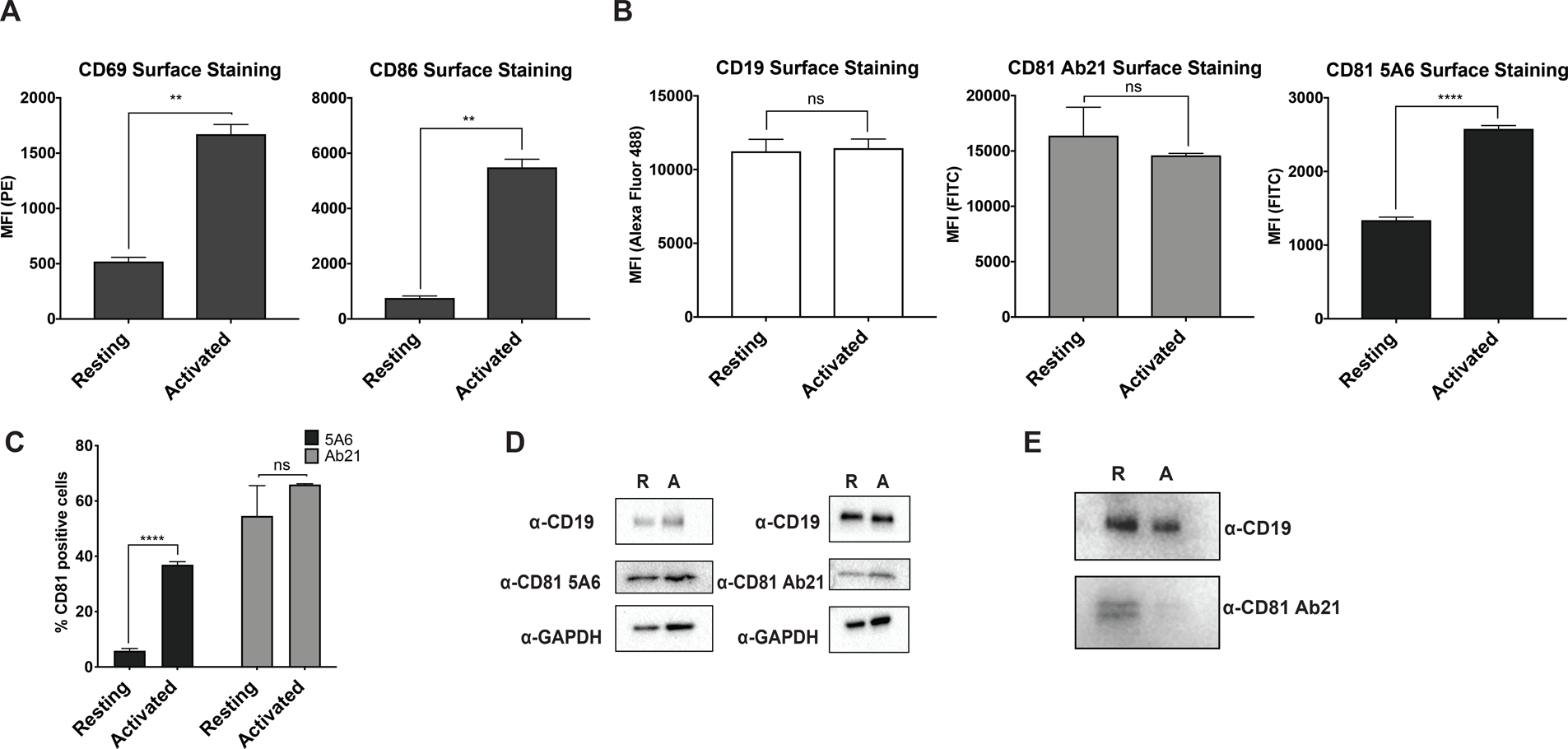
CD81 antibody labeling experiments in resting and activated primary human B cells. **(A)** Surface staining of the B cell activation markers, CD86 and CD69, in resting B cells and cells activated with IgM, IgG Fab’2. **(B)** Surface staining of CD19 and CD81 with antibody 5A6 and Ab21. **(C)** Percent of CD81 positive cells labeled with 5A6 or Ab21. **(D)** Western blots of total protein lysate. “R” represents resting cells and “A” represents activated cells. **(E)** Immunopurification of CD19 from resting and activated primary human B cells, followed by Western blotting for CD19, and for CD81 using Ab21. “R” represents resting cells and “A” represents activated cells. For all panels, data are shown as mean ± SEM. Three replicates were performed for CD81 5A6 and Ab21 staining, and two replicates were performed for CD19, CD69, and CD86 staining conditions. Statistical analysis was performed in GraphPad Prism using an unpaired t test. **p < 0.01; ***p < 0.001, ****p<0.0001.

## DISCUSSION

Tetraspanins control a wide range of physiological processes by interacting with partner proteins (Hemler, 2005), yet there is remarkably little structural or mechanistic information about how tetraspanins bind and regulate their molecular partners. Here, using a combination of molecular engineering, X-ray crystallography, and cell-based assays we investigated the interaction between the prototypical tetraspanin CD81 and its biochemical partner CD19, the key signaling subunit of the B cell co-receptor complex.

Our studies show that the ectodomains of CD19 and CD81 interact dynamically during B cell co-receptor trafficking and signaling upon B cell activation. Our structure of the complex between the 5A6 F_ab_ and the extracellular domain of CD81 reveals that 5A6 binds to an unusual conformational epitope, which is masked in the CD81-CD19 complex. Other CD81 antibodies have epitopes in nearby regions of the CD81 large extracellular loop, yet none rely exclusively on helix C and D for binding or approach CD81 from a similar angle (**Supplementary Figure 3**), suggesting that the exact details of antigen recognition geometry give rise to the unique properties of 5A6. Using immunoprecipitation and flow cytometry, we found that the association between CD19 and CD81 is dynamic and that dissociation of CD19 from CD81 may regulate the association of CD19 with the B cell receptor **(Figure 5)**. This information could be exploited to develop novel co-receptor antibody therapeutics to selectively target activated B cells and to guide development of conformationally selective antibodies targeting other therapeutically relevant tetraspanin-partner protein complexes. Indeed, the 5A6 antibody itself has recently shown promise as a therapeutic lead, since it can selectively target malignant B cells while sparing normal cells (Vences-Catalán et al., 2019).

**Figure 5:**
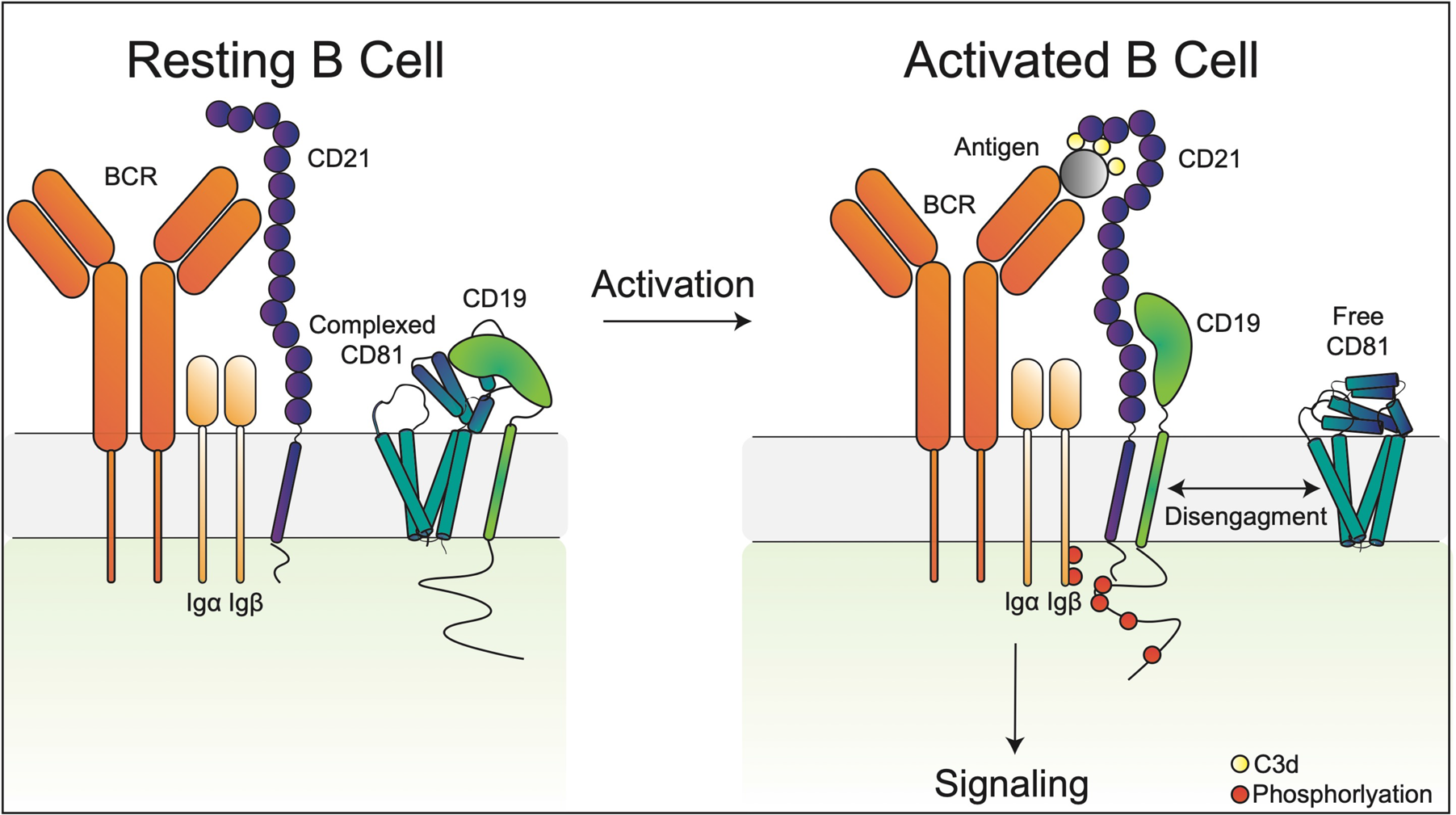
Proposed model for the disengagement of the CD81 during B cell activation. Upon B cell activation, dissociation of the B cell co-receptor complex allows CD19 to freely diffuse in the membrane and interact with the BCR, leading to amplified signaling through the BCR and activation of the B cell.

Our observation of regulated dissociation of the CD19-CD81 complex is particularly interesting in view of prior experiments suggesting that CD81 may regulate the diffusion of CD19 by immobilizing it in distinct locations in the membrane (Cherukuri et al., 2004; Mattila et al., 2013). For example, super resolution microscopy has shown that in resting B cells, the mobility of a large proportion of CD19 is very low, while in CD81-deficient B cells CD19 diffusion shifts strongly to a faster moving population (Mattila et al., 2013). This control of CD19 diffusion by CD81 could allow regulation of CD19’s interaction with the BCR, for example, to prevent high-level constitutive signaling (Mattila et al., 2013). Upon B cell activation, dissociation of the CD19-CD81 complex allows CD19 to freely diffuse in the membrane and interact with the BCR, leading to amplified signaling through the BCR and activation of the B cell **(Figure 5)**. During B cell activation, free CD81 could also become involved in other aspects of B cell biology. For example, others have shown that CD81 redistributes to the immune synapse of activated B cells (Mittelbrunn et al., 2002). Several integrins have been reported to associate with CD81 and are involved in CD81-mediated adhesion in the immune synapse in activated B cells as well as in B cell trafficking to lymphoid organs (Levy et al., 1998).

Beyond our finding that the ectodomains of CD19 and CD81 interact to control receptor trafficking and localization, there are other lines of evidence suggesting that the use of the large extracellular loop to bind partner proteins will be a general property of the tetraspanin protein family. First, the transmembrane regions of tetraspanins are highly conserved, but the large extracellular loop varies among family members in both size and sequence. Sequence analysis reveals that within the large extracellular loop, tetraspanins contain a low-polarity hypervariable region that may be involved in tetraspanin binding partner recognition (Stipp et al., 2003). Additionally, it is known that several other tetraspanins and their binding partners, such ADAM10 and the C8 family of tetraspanins and the tetraspanin CD151 and integrin α_3_β_1_, rely on their ectodomains for binding of partner proteins (Noy et al., 2016; Yauch et al., 2000). Like the CD19-CD81 complex, other tetraspanin-partner protein complexes may also be dynamically regulated upon changes in cell state, and conformationally specific antibodies may serve as powerful tools to investigate and control tetraspanin biology.

## ACKNOWLEDGMENTS

We would like to thank Izabela Durzynska for cloning of CD9/CD81 chimeras, the CD19 transmembrane domain chimera, and cloning of the guide RNA. We would also like to thank Sanchez Jarrett for assistance with harvesting crystals and Megan Sjodt-Gable for input on figure design. Financial support for this work was provided by NIH grants R35 CA220340 (S.C.B.), F31 HL147459 (K.J.S.) and DP5 OD02134 (A.C.K.).

## AUTHOR CONTRIBUTIONS

The overall project was designed and developed by K.J.S., A.C.K., and S.C.B. Molecular cloning, protein purification, flow cytometry, and primary B cell experiments were conducted by K.J.S. Crystallization and data collection was performed by K.J.S. with guidance from T.C.M.S. Data processing and structure refinement was carried out K.J.S. with input from T.C.M.S, A.C.K, and S.C.B. The manuscript was written by K.J.S., A.C.K., and S.C.B. with input from T.C.M.S.

## STAR METHODS

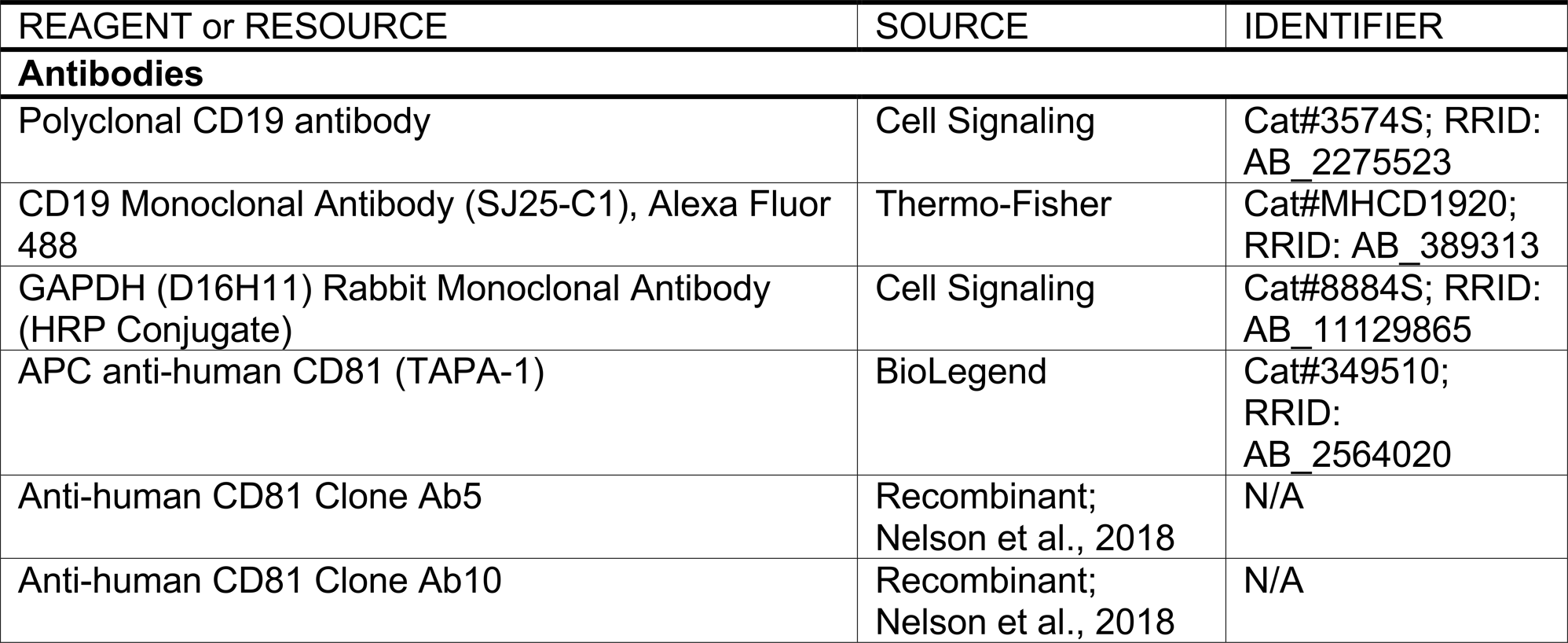
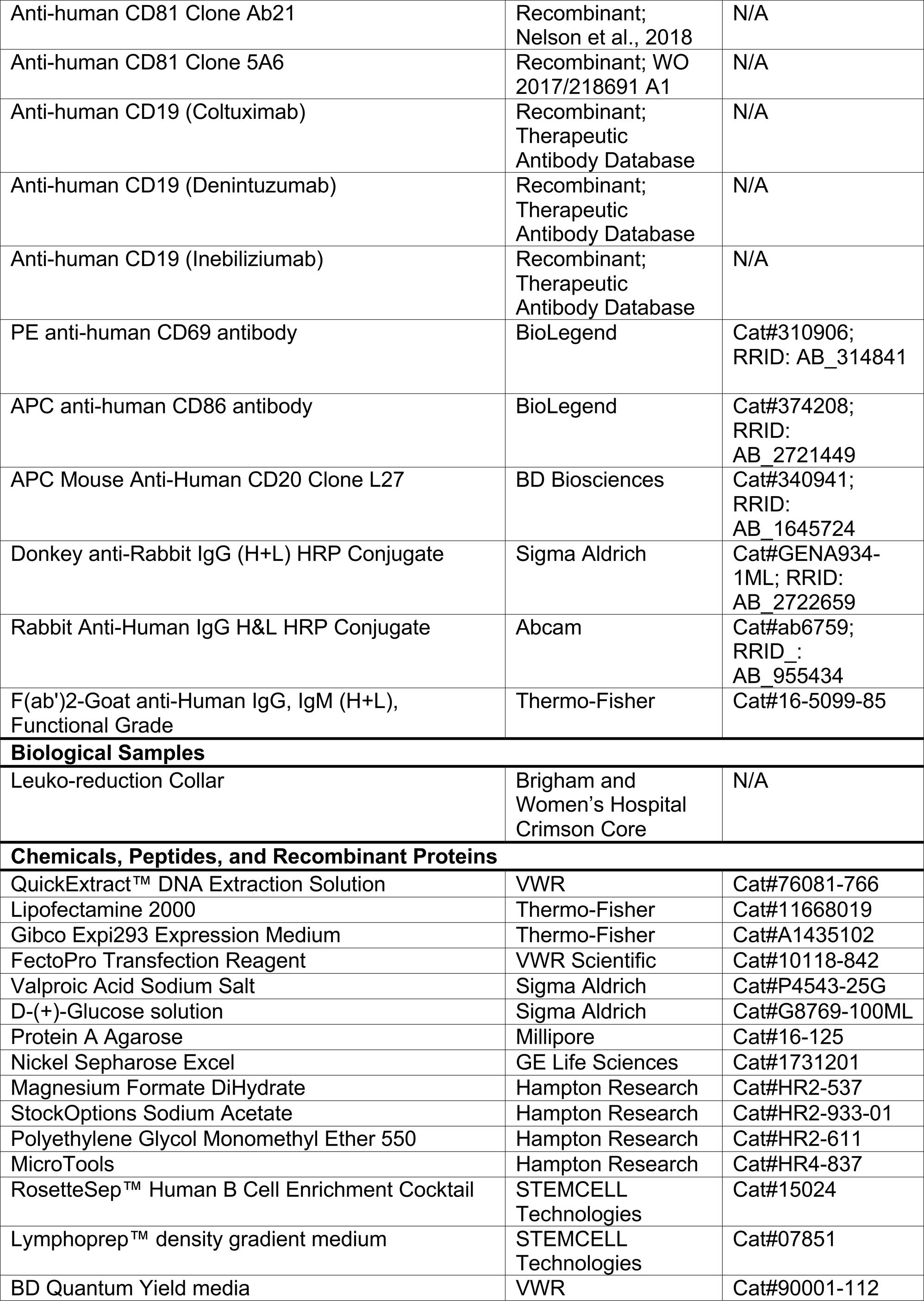
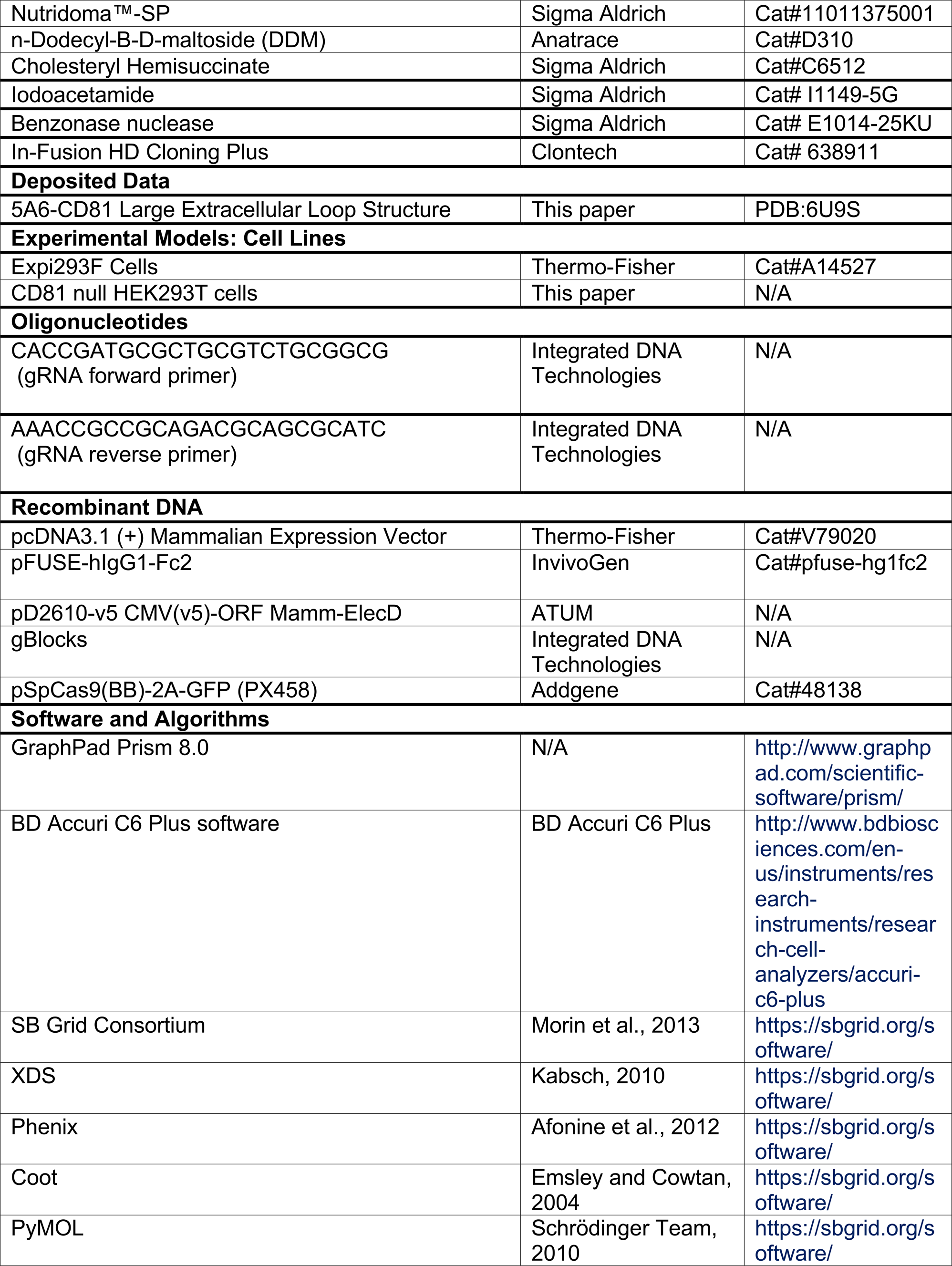

## METHOD DETAILS

### CD81 Knockout HEK293T Cell Line

To generate a CD81^−/−^ HEK293T cell line, the CHOPCHOP guide design server (http://chopchop.cbu.uib.no/) was used to select guide sequences targeting exon 1 of the *CD81* gene. The two complementary DNA strands of the guide sequences (IDT Technologies) were annealed in 10 mM Tris pH 8.0, 50 mM NaCl, 1 mM EDTA and then subcloned into a pSpCas9 WT-2A-GFP vector. The resulting pSpCas9 WT-2A-GFP cDNA was transfected into HEK293T cells using polyethyleneimine. Cells expressing GFP were sorted into 96-well plates by flow cytometry 48 hours after transfection. Clonal populations were allowed to expand for 4 weeks. Genomic DNA was extracted from individual clones, and the CD81 gene was amplified by PCR and sequenced to confirm the presence of targeted mutations. The loss of CD81 expression was confirmed by flow cytometry.

### CD19 Export Assay

CD81^−/−^ HEK293T cells were seeded at 100,000 cells/well in 24 well plates 12-18 hours prior to transfection. CD81^−/−^ HEK293T cells were transfected using Lipofectamine 2000 with either with either 1.5 μg of empty pcDNA3.1(+) vector, 0.75 μg of CD19 DNA and 0.75 μg of empty pcDNA3.1(+) vector DNA (CD19 condition), 0.75 μg of CD19 DNA and 0.75 μg of CD81 DNA (CD19+CD81 condition), or 0.75 μg of CD19 DNA and 0.75 μg of a CD81 chimera DNA. 36-48 hours after transfection, cells were harvested in phosphate buffered saline (PBS) supplemented with 3 mM EDTA, transferred to a 96 well V-bottom plate, and then washed twice with PBS. Cells were then incubated on ice for 20 minutes with 2 μg/mL Alexa 488-anti-CD19 (ThermoFisher) and APC-anti-CD81 (BioLegend) in 20 mM HEPES buffer pH 7.4, containing 150 mM NaCl, and 0.1% BSA. Cells were washed two times with PBS and analyzed on a BD Accuri C6 flow cytometer.

### Cloning of Constructs

#### CD19-CD81 Fusion Protein

The CD19-CD81 fusion was cloned into pcDNA3.1(+) with an N-terminal haemagglutinin signal sequence followed by a FLAG epitope tag and a 3C protease cleavage site. Residues 20-329 of CD19 (ectodomain, transmembrane domain, and first 15 cytoplasmic amino acids) were connected to full length CD81 using a GGSG linker.

#### CD81 Chimeras

CD81 chimeras were constructed by PCR and subcloned into pcDNA3.1(+). All chimeras were created within the backbones of wild-type human CD9, *C.elegans* Tspan15, or human claudin-4. The following domain boundaries were used:

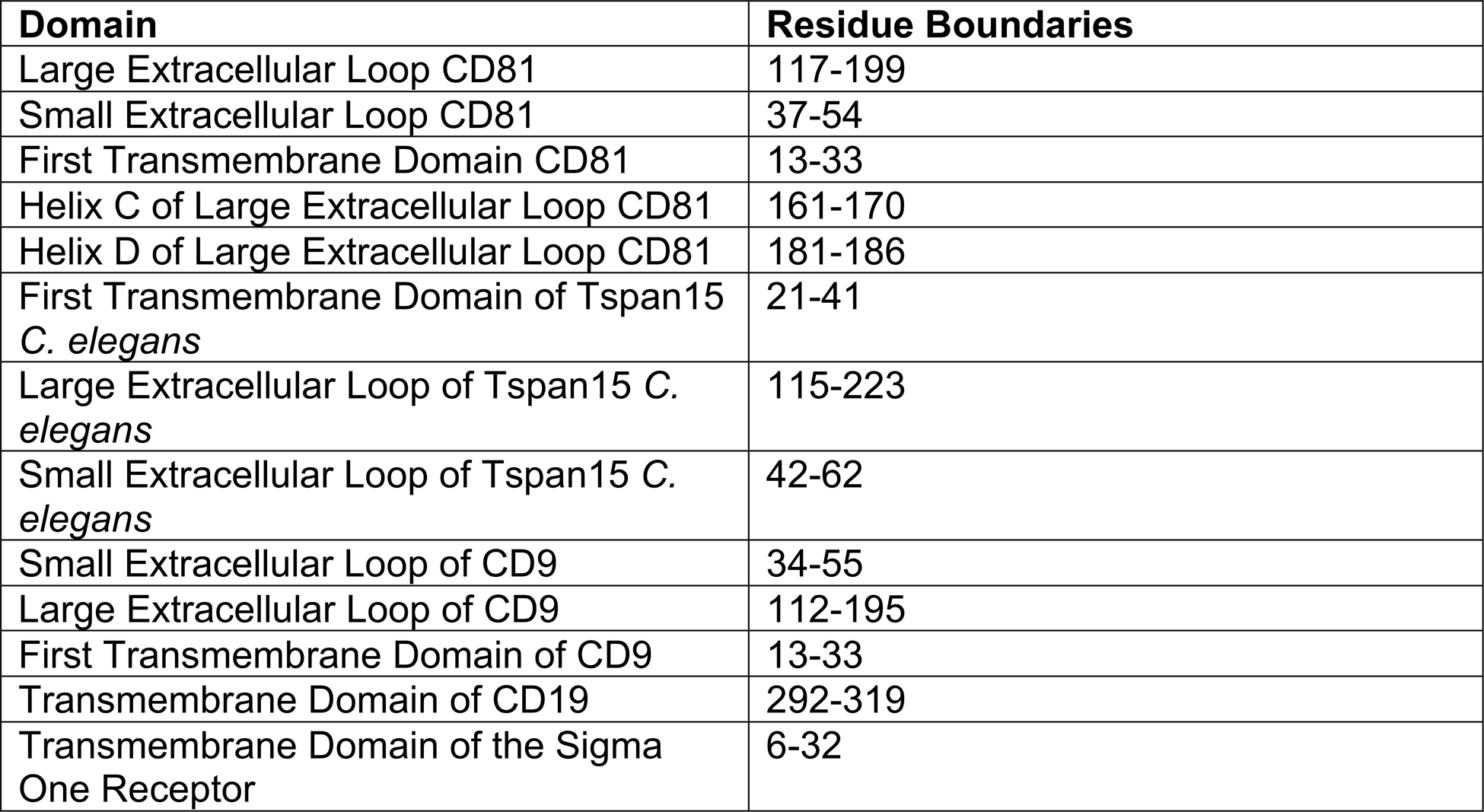

#### Antibodies 5A6, Ab5, Ab10, Ab21, Denintuzumab, Coltuximab, and Inebilizumab

The variable regions of each antibody heavy chain were subcloned into the pFUSE-hIgG1-Fc2 vector (Invitrogen). The variable region of the light chains and the human kappa constant sequence with an N terminal “MDWTWRILFLVAAATGAHS” signal sequence were cloned in the pD2610-v5 vector (ATUM). An additional construct of the 5A6 antibody was also cloned, with a 3C protease site flanked by a Gly-Gly-Ser-Gly linker inserted into the hinge region of the heavy chain, allowing for generation of the 5A6 Fab after cleavage with 3C protease for use in crystallography.

### Validation of CD19-CD81 Fusion in CD19 Export Assay

CD81^−/−^ HEK293T cells were seeded at 100,000 cells/well in 24 well plates 12-18 hours prior to transfection. CD81^−/−^ HEK293T cells were transfected using Lipofectamine 2000 with either 0.75 μg of CD19 DNA and 0.75 μg of empty vector DNA (CD19 condition), 0.75 μg of CD19 DNA and 0.75 μg of CD81 DNA (CD19+CD81 condition), or 0.75 μg of CD19-CD81 DNA and 0.75 μg empty vector DNA (CD19-CD81 fusion protein condition). 48 hours after transfection, cells were harvested in PBS containing 3 mM EDTA, transferred to a 96 well V-bottom plate, and then washed two times with PBS. Cells were then incubated on ice for 20 minutes with 2 μg/mL Alexa 488-anti-CD19 (ThermoFisher), Coltuximab (recombinant), Denintuzumab (recombinant), or Inebilizumab (recombinant) in 20 mM HEPES buffer pH 7.4, 150 mM NaCl, and 0.1% BSA. Cells stained with recombinant CD19 antibodies were detected with Goat anti Human IgG (H+L) Secondary Antibody, Alexa Fluor 488 (Invitrogen) at 1 μg/mL. Cells were washed two times with PBS and analyzed on a BD Accuri C6 flow cytometer.

### Expression and Purification of 5A6 Fab-CD81 LEL Complex

The heavy and light chains of the 5A6 antibody and the CD81 large extracellular loop were co-expressed in Expi293F cells. 600 mL of Expi293F cells maintained in Expi293 expression media were grown to a density of 2.8 × 10^6^ cells/mL and then transiently transfected with heavy chain 5A6, light chain 5A6, and CD81 LEL DNA (0.48 mg total DNA) and FectoPro transfection reagent (Polyplus) at a 1:1 DNA/FectoPro ratio. 20 hours after transfection, the cells were fed 5 mM Valproic acid sodium salt (Sigma-Aldrich) and 5.5 mL of 45% D-(+)-Glucose solution (Sigma-Aldrich). Transfected cells were cultured for 7 days to produce protein and then the media was collected and separated from the cells by centrifugation at 4000g for 15 minutes at 4°C. The cultured media was loaded onto protein A resin (Millipore). The resin was washed with 100 mL 20 mM Tris buffer pH 7.4, containing 150 mM NaCl and then bound protein was eluted in 20 mL 100 mM glycine buffer pH 3.0. Elution fractions were immediately neutralized with 1M Tris buffer pH 7.5. Eluted protein was buffer exchanged into 20 mM Tris buffer pH 7.4, containing 150 mM NaCl and then 3C protease was added at a 1:1 w/w ratio and incubated overnight at 4°C. The purity of the eluted protein and efficiency of cleavage was assessed on an SDS-PAGE Coomassie-stained gel.

The cleaved, recovered 5A6 Fab and CD81 LEL was then applied to nickel resin to select for Fab bound to CD81. Nickel resin was washed with 20 mM Tris buffer pH 7.4, containing 150 mM NaCl and 20 mM imidazole pH 7.4 and then eluted in the same buffer with 350 mM imidazole. The sample was then applied to Protein A resin to remove any residual free Fc. Flow through from the protein A resin was then concentrated with a centrifugal filter, and the purified 5A6 Fab-LEL complex was isolated on an S200 size exclusion column in 20 mM Tris buffer pH 7.4, containing 150 mM NaCl. The purity of fractions corresponding to the 5A6 Fab-LEL complex peak were assessed on an SDS-PAGE Coomassie-stained gel and then pooled and concentrated to 7.2 mg/mL for crystallography. The concentrated complex was stored at 4°C for ~36 hours prior to setting trays.

### Crystallization of 5A6 Fab-CD81 LEL Complex

Crystals of the 5A6 Fab-CD81 LEL complex were grown in 96-well sitting drops at room temperature. Branched crystals of the complex (7.5 mg/mL) grew after 48 hours in 0.2 M magnesium formate dihydrate, 0.1 M sodium acetate trihydrate pH 4.0, 18% w/v polyethylene glycol monomethyl ether 5,000. Crystals harvested for data collection were grown in 100 nL drops (50 nL protein + 50 nL precipitant) with a 50 μL reservoir solution. To isolate single crystal fragments, crystals were cut with MicroTools (Hampton Research). Crystals were cryoprotected by supplementing the mother liquor with 20% glycerol (v/v). Individual crystals were flash frozen in liquid nitrogen and stored until data collection.

Data collection was performed at Advanced Photon Source NE-CAT beamline 24 ID-C. Diffraction images were processed and scaled using XDS (Kabsch, 2010). The crystals were indexed in the space group P2_1_2_1_2_1_ with unit-cell dimensions of 40.0, 96.9, 297.1 Å. Data to a maximum resolution of 2.4 Å were used for structure solution and refinement. The structure was solved by molecular replacement using the program Phaser (McCoy et al., 2007). Search models included the heavy and light chain (chain H and L) from PDB entry 4S1D and residues 112-201 of chain A (large extracellular loop of CD81) from PDB entry 5TCX. Two copies of the Fab-CD81 complex were modeled in the asymmetric unit. The structural model was built by auto-build and iterative cycles of manual rebuilding and refinement in Coot and PHENIX (Afonine et al., 2012).

### Expression and Purification of Ab5, Ab10, Ab21, and 5A6 for use in Flow Cytometry

Expi293F cells maintained in Expi293 expression media were grown to cell density of 2.8 × 10^6^ cells/mL and then transiently transfected. For each antibody, plasmids encoding the heavy and light chain were transfected at a 1:2 molar ratio (0.8 mg total DNA/liter cells) with FectoPro transfection reagent (Polyplus) at 1:1 DNA/FectoPro ratio. 20-24 hours after transfection, the cells were fed with 5 mM Valproic acid sodium salt (Sigma-Aldrich) and 45% D-(+)-Glucose solution (Sigma-Aldrich). 4-7 days after transfection, the media was collected and separated from the cells by centrifugation at 4000g for 15 minutes at 4 °C. Cultured media was loaded onto protein A resin (Millipore). The resin was then washed with 50 mL of 20 mM HEPES buffer pH 7.4, containing 150 mM NaCl and then bound protein was eluted in 10 mL of 100 mM glycine buffer, pH 3.0. Elution fractions were immediately neutralized with 1 M HEPES buffer pH 7.4. Eluted protein was buffer exchanged into 20 mM HEPES buffer pH 7.4, containing 150 mM NaCl.

### Isolation of Primary Human B Cells

A leuko-reduction collar was obtained from the Brigham and Women’s Hospital Crimson Core with patient information deidentified. All methods were carried out in accordance with relevant guidelines and regulations. All experimental protocols were reviewed and approved as exempt by the Harvard Faculty of Medicine Institutional Review Board. Primary human B cells were isolated from fresh leuko-reduction collar blood by using 750 μL of RosetteSep™ Human B Cell Enrichment Cocktail (Stemcell Technologies) for 10 mL of collar blood. The RosetteSep™ cocktail was incubated with collar blood for 20 minutes at room temperature, and then 10 mL of PBS supplemented with 2% FBS was added to the collar blood and mixed gently. The diluted collar blood was then layered on top of 10 mL Lymphoprep™ density gradient medium (Stemcell Technologies). After centrifugation at 1200 g for 20 minutes, the mononuclear cell layer was harvested and washed twice with PBS supplemented with 2% FBS. Purified cells were then resuspended in warm BD Quantum Yield media (BD Biosciences) supplemented with 10% FBS and 1:50 Nutridoma™-SP (Sigma Aldrich) at 10^6^ cells/mL.

### B Cell Activation Assay

Purified cells were resuspended in warm BD Quantum Yield media supplemented with 10% FBS and 1:50 Nutridoma™-SP (Sigma Aldrich) at 10^6^ cells/mL and allowed to rest for 30 minutes at 37°C and 5% CO_2_. Cells were then pipeted several times to disperse clumps and were split into 96 wells for stimulation assays (100 μL per well). Cells were allowed to rest for 1 hour at 37°C and 5% CO_2_. Either 100 μL of media (unstimulated condition) or 100 μL of media containing 20 μg/mL F(ab’)2-Goat anti-Human IgG, IgM (H+L) (Invitrogen) was added to each well. 72 hours later, cells were harvested at 500 g for 5 min, washed once with PBS, stained with 2 μg/mL CD69, CD86, CD81, or CD19 antibody in 20mM HEPES buffer (pH 7.4), 150 mM NaCl, and 0.1% BSA for 20 minutes on ice, washed twice with PBS, and analyzed on a BD Accuri C6 flow cytometer.

### Western Blot Analysis of Total CD19 and CD81 Protein Abundance in Resting and Activated Primary Human B Cells

Cell lysates were run on a non-reducing SDS-PAGE gel, and the protein was transferred to a nitrocellulose membrane. The membrane was then gently rocked in in TBST (0.1% Tween-20 in Tris-buffered-saline) blocking buffer with 5% (w/v) non-fat milk powder at room temperature for 1.5 hours. The blocked membrane was cut and then incubated in TBST with 1% non-fat milk powder containing either GADPH-HRP conjugate antibody (Cell Signaling; 1:10,000 dilution), CD19 SJ25 antibody (Cell Signaling; 1:500 dilution) or CD81 5A6 antibody (Recombinant, 0.4 mg/mL; 1:100 dilution). Antibody incubations were performed overnight at 4 °C with shaking. The CD19 blot was incubated with Donkey anti Rabbit IgG (H+L) HRP Conjugate (Thermo Fischer) diluted 1:5,000 in TBST containing 1% non-fat milk powder. The CD81 blot was incubated with Rabbit Anti-Human IgG H&L HRP Conjugate (Abcam) diluted 1:5,000 in TBST with 1% non-fat milk powder. Secondary antibody incubations were performed for 1 h at room temperature with shaking. Prior to chemiluminescent detection, blots were washed with TBST three times for 10 minutes each time. Western blots were developed with Western Lightning® Plus-ECL, Enhanced Chemiluminescence Detection Kit (PerkinElmer).

### CD19-CD81 Pulldown and Western Blots in Resting and Activated Primary Human B Cells

Purified cells were resuspended in warm BD Quantum Yield media supplemented with 10% FBS and 1:50 Nutridoma™-SP (Sigma Aldrich) at 1 million cells/mL and allowed to rest for 30 minutes at 37 °C and 5% CO_2_. Cells were then pipeted several times to disperse clumps and were split into a 6 well plate for stimulation (1 mL per well). Cells were allowed to rest for 1 hour at 37 °C and 5% CO_2_. Either 1 mL of media (unstimulated condition) or 1 mL of media containing 10 μg/mL F(ab’)2-Goat anti-Human IgG, IgM (H+L) (Invitrogen) was added to each well. 72 hours later, cells were harvested at 500 g for 10 min. Cells were lysed in 50 μL 20 mM HEPES buffer pH 7.4, containing 2 mM MgCl_2_, 2 mg/mL iodoacetamide and 1:100,000 v:v benzonase nuclease. Lysate was centrifuged at 16,000 g for 15 min, and then membranes were resuspended in 200 μL 20 mM HEPES buffer pH 7.4, containing 1% n-Dodecyl-B-D-maltoside (DDM) (Anatrace), 0.1% cholesteryl hemisuccinate (Sigma Aldrich), 250 mM NaCl, and 10% v/v glycerol and then incubated at 4°C with rotating for 2 hours. 200 μL of solubilized protein was then applied to 10 μL protein A resin preincubated with 15 μg Coltuximab (anti-CD19, recombinant) and incubated at 4°C with rotating for 2 hours. Samples were centrifuged at 200g for 1 minute and washed twice with 1 mL of 20 mM HEPES buffer pH 7.4, containing 0.1% n-Dodecyl-B-D-maltoside (DDM) (Anatrace), 0.01% cholesteryl hemisuccinate (Sigma Aldrich), 250 mM NaCl, and 1% v/v glycerol. Samples were eluted in 50 μL 2X SDS loading dye and separated by SDS-PAGE under non-reducing conditions. The membrane was blocked with 5% (w/v) non-fat milk powder in TBST (0.1% Tween-20 in Tris-buffered-saline) at room temperature for 1.5 hours. The blocked membrane was cut and then incubated with shaking overnight at 4°C with either CD19 SJ25 antibody (Cell Signaling; 1:1000 dilution) or CD81 Ab21 antibody (Recombinant, 1 mg/mL; 1:1000 dilution) in TBST containing 1% non-fat milk powder. The CD19 blot was incubated with donkey anti Rabbit IgG (H+L) HRP Conjugate (Thermo Fischer) diluted 1:5,000 in TBST containing 1% non-fat milk powder. The CD81 blot was incubated with Rabbit Anti-Human IgG H&L HRP Conjugate (Abcam) diluted 1:5,000 in TBST with 1% non-fat milk powder. Secondary antibody incubations were performed for 1 h at room temperature with shaking. Prior to chemiluminescent detection, blots were washed with TBST three times for 10 minutes each time. Western blots were developed with Western Lightning® Plus-ECL, Enhanced Chemiluminescence Detection Kit (PerkinElmer).

## REFERENCES

Afonine, P.V., Grosse-Kunstleve, R.W., Echols, N., Headd, J.J., Moriarty, N.W., Mustyakimov, M., Terwilliger, T.C., Urzhumtsev, A., Zwart, P.H., and Adams, P.D. (2012). Towards automated crystallographic structure refinement with phenix.refine. Acta Crystallogr. D Biol. Crystallogr. 68, 352–367.

Barrena, S., Almeida, J., Yunta, M., López, A., Fernández-Mosteirín, N., Giralt, M., Romero, M., Perdiguer, L., Delgado, M., Orfao, A., et al. (2005). Aberrant expression of tetraspanin molecules in B-cell chronic lymphoproliferative disorders and its correlation with normal B-cell maturation. Leukemia 19, 1376–1383.

Berditchevski, F., and Odintsova, E. (2007). Tetraspanins as Regulators of Protein Trafficking. Traffic 8, 89–96.

Bradbury, L.E., Kansas, G.S., Levy, S., Evans, R.L., and Tedder, T.F. (1992). The CD19/CD21 signal transducing complex of human B lymphocytes includes the target of antiproliferative antibody-1 and Leu-13 molecules. J. Immunol. 149, 2841–2850.

Braig, F., Brandt, A., Goebeler, M., Tony, H.-P., Kurze, A.-K., Nollau, P., Bumm, T., Böttcher, S., Bargou, R.C., and Binder, M. (2016). Resistance to anti-CD19/CD3 BiTE in acute lymphoblastic leukemia may be mediated by disrupted CD19 membrane trafficking. Blood blood-2016-05-718395.

Brentjens, R.J., Davila, M.L., Riviere, I., Park, J., Wang, X., Cowell, L.G., Bartido, S., Stefanski, J., Taylor, C., Olszewska, M., et al. (2013). CD19-Targeted T Cells Rapidly Induce Molecular Remissions in Adults with Chemotherapy-Refractory Acute Lymphoblastic Leukemia. Sci. Transl. Med. 5, 177ra38–177ra38.

Carter, R.H., and Barrington, R.A. (2004). Signaling by the CD19/CD21 complex on B cells. Curr. Dir. Autoimmun. 7, 4–32.

Carter, R.H., and Fearon, D.T. (1992). CD19: Lowering the Threshold for Antigen Receptor Stimulation of B Lymphocytes. Science 256, 105–107.

Cherukuri, A., Shoham, T., Sohn, H.W., Levy, S., Brooks, S., Carter, R., and Pierce, S.K. (2004). The Tetraspanin CD81 Is Necessary for Partitioning of Coligated CD19/CD21-B Cell Antigen Receptor Complexes into Signaling-Active Lipid Rafts. J. Immunol. 172, 370–380.

Gauld, S.B., Porto, J.M.D., and Cambier, J.C. (2002). B Cell Antigen Receptor Signaling: Roles in Cell Development and Disease. Science 296, 1641–1642.

Grupp, S.A., Kalos, M., Barrett, D., Aplenc, R., Porter, D.L., Rheingold, S.R., Teachey, D.T., Chew, A., Hauck, B., Wright, J.F., et al. (2013). Chimeric antigen receptor-modified T cells for acute lymphoid leukemia. N. Engl. J. Med. 368, 1509–1518.

Hemler, M.E. (2005). Tetraspanin functions and associated microdomains. Nat. Rev. Mol. Cell Biol. 6, 801–811.

Hemler, M.E. (2008). Targeting of tetraspanin proteins — potential benefits and strategies. Nat. Rev. Drug Discov. 7, 747.

Kabsch, W. (2010). XDS. Acta Crystallogr. D Biol. Crystallogr. 66, 125–132.

Kalos, M., Levine, B.L., Porter, D.L., Katz, S., Grupp, S.A., Bagg, A., and June, C.H. (2011). T cells with chimeric antigen receptors have potent antitumor effects and can establish memory in patients with advanced leukemia. Sci. Transl. Med. 3, 95ra73.

Kochenderfer, J.N., Dudley, M.E., Feldman, S.A., Wilson, W.H., Spaner, D.E., Maric, I., Stetler-Stevenson, M., Phan, G.Q., Hughes, M.S., Sherry, R.M., et al. (2012). B-cell depletion and remissions of malignancy along with cytokine-associated toxicity in a clinical trial of anti-CD19 chimeric-antigen-receptor–transduced T cells. Blood 119, 2709–2720.

Levy, S., Todd, S.C., and Maecker, H.T. (1998). CD81 (TAPA-1): a molecule involved in signal transduction and cell adhesion in the immune system. Annu. Rev. Immunol. 16, 89–109.

Levy, S., Marabelle, A., Rajapaksa, R., Vences-catalán, F., Kuo, C.-C., Liu, J., and Levy, R. (2017). Humanized and chimeric monoclonal antibodies to cd81.

Maecker, H.T., and Levy, S. (1997). Normal lymphocyte development but delayed humoral immune response in CD81-null mice. J. Exp. Med. 185, 1505–1510.

Matsumoto, A.K., Martin, D.R., Carter, R.H., Klickstein, L.B., Ahearn, J.M., and Fearon, D.T. (1993). Functional dissection of the CD21/CD19/TAPA-1/Leu-13 complex of B lymphocytes. J. Exp. Med. 178, 1407–1417.

Mattila, P.K., Feest, C., Depoil, D., Treanor, B., Montaner, B., Otipoby, K.L., Carter, R., Justement, L.B., Bruckbauer, A., and Batista, F.D. (2013). The actin and tetraspanin networks organize receptor nanoclusters to regulate B cell receptor-mediated signaling. Immunity 38, 461–474.

McCoy, A.J., Grosse-Kunstleve, R.W., Adams, P.D., Winn, M.D., Storoni, L.C., and Read, R.J. (2007). Phaser crystallographic software. J. Appl. Crystallogr. 40, 658–674.

Mei, H.E., Schmidt, S., and Dörner, T. (2012). Rationale of anti-CD19 immunotherapy: an option to target autoreactive plasma cells in autoimmunity. Arthritis Res. Ther. 14, S1.

Mittelbrunn, M., Yáñez-Mó, M., Sancho, D., Ursa, Á., and Sánchez-Madrid, F. (2002). Cutting Edge: Dynamic Redistribution of Tetraspanin CD81 at the Central Zone of the Immune Synapse in Both T Lymphocytes and APC. J. Immunol. 169, 6691–6695.

Miyazaki, T., Müller, U., and Campbell, K.S. (1997). Normal development but differentially altered proliferative responses of lymphocytes in mice lacking CD81. EMBO J. 16, 4217–4225.

Nelson, B., Adams, J., Kuglstatter, A., Li, Z., Harris, S.F., Liu, Y., Bohini, S., Ma, H., Klumpp, K., Gao, J., et al. (2018). Structure-Guided Combinatorial Engineering Facilitates Affinity and Specificity Optimization of Anti-CD81 Antibodies. J. Mol. Biol. 430, 2139–2152.

Noy, P.J., Yang, J., Reyat, J.S., Matthews, A.L., Charlton, A.E., Furmston, J., Rogers, D.A., Rainger, G.E., and Tomlinson, M.G. (2016). TspanC8 Tetraspanins and A Disintegrin and Metalloprotease 10 (ADAM10) Interact via Their Extracellular Regions EVIDENCE FOR DISTINCT BINDING MECHANISMS FOR DIFFERENT TspanC8 PROTEINS. J. Biol. Chem. 291, 3145–3157.

Oren, R., Takahashi, S., Doss, C., Levy, R., and Levy, S. (1990). TAPA-1, the target of an antiproliferative antibody, defines a new family of transmembrane proteins. Mol. Cell. Biol. 10, 4007–4015.

Porter, D.L., Levine, B.L., Kalos, M., Bagg, A., and June, C.H. (2011). Chimeric antigen receptor-modified T cells in chronic lymphoid leukemia. N. Engl. J. Med. 365, 725–733.

Rajesh, S., Sridhar, P., Tews, B.A., Fénéant, L., Cocquerel, L., Ward, D.G., Berditchevski, F., and Overduin, M. (2012). Structural Basis of Ligand Interactions of the Large Extracellular Domain of Tetraspanin CD81. J. Virol. 86, 9606–9616.

Schmidt, H.R., Zheng, S., Gurpinar, E., Koehl, A., Manglik, A., and Kruse, A.C. (2016a). Crystal structure of the human σ1 receptor. Nature 532, 527–530.

Schmidt, T.H., Homsi, Y., and Lang, T. (2016b). Oligomerization of the Tetraspanin CD81 via the Flexibility of Its δ-Loop. Biophys. J. 110, 2463–2474.

Shoham, T., Rajapaksa, R., Boucheix, C., Rubinstein, E., Poe, J.C., Tedder, T.F., and Levy, S. (2003). The Tetraspanin CD81 Regulates the Expression of CD19 During B Cell Development in a Postendoplasmic Reticulum Compartment. J. Immunol. 171, 4062–4072.

Shoham, T., Rajapaksa, R., Kuo, C.-C., Haimovich, J., and Levy, S. (2006). Building of the Tetraspanin Web: Distinct Structural Domains of CD81 Function in Different Cellular Compartments. Mol. Cell. Biol. 26, 1373–1385.

Stipp, C.S., Kolesnikova, T.V., and Hemler, M.E. (2003). Functional domains in tetraspanin proteins. Trends Biochem. Sci. 28, 106–112.

Tsitsikov, E.N., Gutierrez-Ramos, J.C., and Geha, R.S. (1997). Impaired CD19 expression and signaling, enhanced antibody response to type II T independent antigen and reduction of B-1 cells in CD81-deficient mice. Proc. Natl. Acad. Sci. U. S. A. 94, 10844–10849.

Vences-Catalán, F., Kuo, C.-C., Rajapaksa, R., Duault, C., Andor, N., Czerwinski, D.K., Levy, R., and Levy, S. (2019). CD81 is a novel immunotherapeutic target for B cell lymphoma. J. Exp. Med. 216, 1497–1508.

Yauch, R.L., Kazarov, A.R., Desai, B., Lee, R.T., and Hemler, M.E. (2000). Direct Extracellular Contact between Integrin α3β1 and TM4SF Protein CD151. J. Biol. Chem. 275, 9230–9238.

Yazawa, N., Hamaguchi, Y., Poe, J.C., and Tedder, T.F. (2005). Immunotherapy using unconjugated CD19 monoclonal antibodies in animal models for B lymphocyte malignancies and autoimmune disease. Proc. Natl. Acad. Sci. U. S. A. 102, 15178–15183.

van Zelm, M.C., Reisli, I., van der Burg, M., Castaño, D., van Noesel, C.J.M., van Tol, M.J.D., Woellner, C., Grimbacher, B., Patiño, P.J., van Dongen, J.J.M., et al. (2006). An antibody-deficiency syndrome due to mutations in the CD19 gene. N. Engl. J. Med. 354, 1901–1912.

van Zelm, M.C., Smet, J., Adams, B., Mascart, F., Schandené, L., Janssen, F., Ferster, A., Kuo, C.-C., Levy, S., van Dongen, J.J.M., et al. (2010). CD81 gene defect in humans disrupts CD19 complex formation and leads to antibody deficiency. J. Clin. Invest. 120, 1265–1274.

Zuidscherwoude, M., Göttfert, F., Dunlock, V.M.E., Figdor, C.G., van den Bogaart, G., and van Spriel, A.B. (2015). The tetraspanin web revisited by super-resolution microscopy. Sci. Rep. 5, 12201.

