## Supplementary Information for "A dynamic interaction between CD19 and the tetraspanin CD81 controls B cell co-receptor trafficking"

### SUPPLEMENTAL FIGURES AND LEGENDS

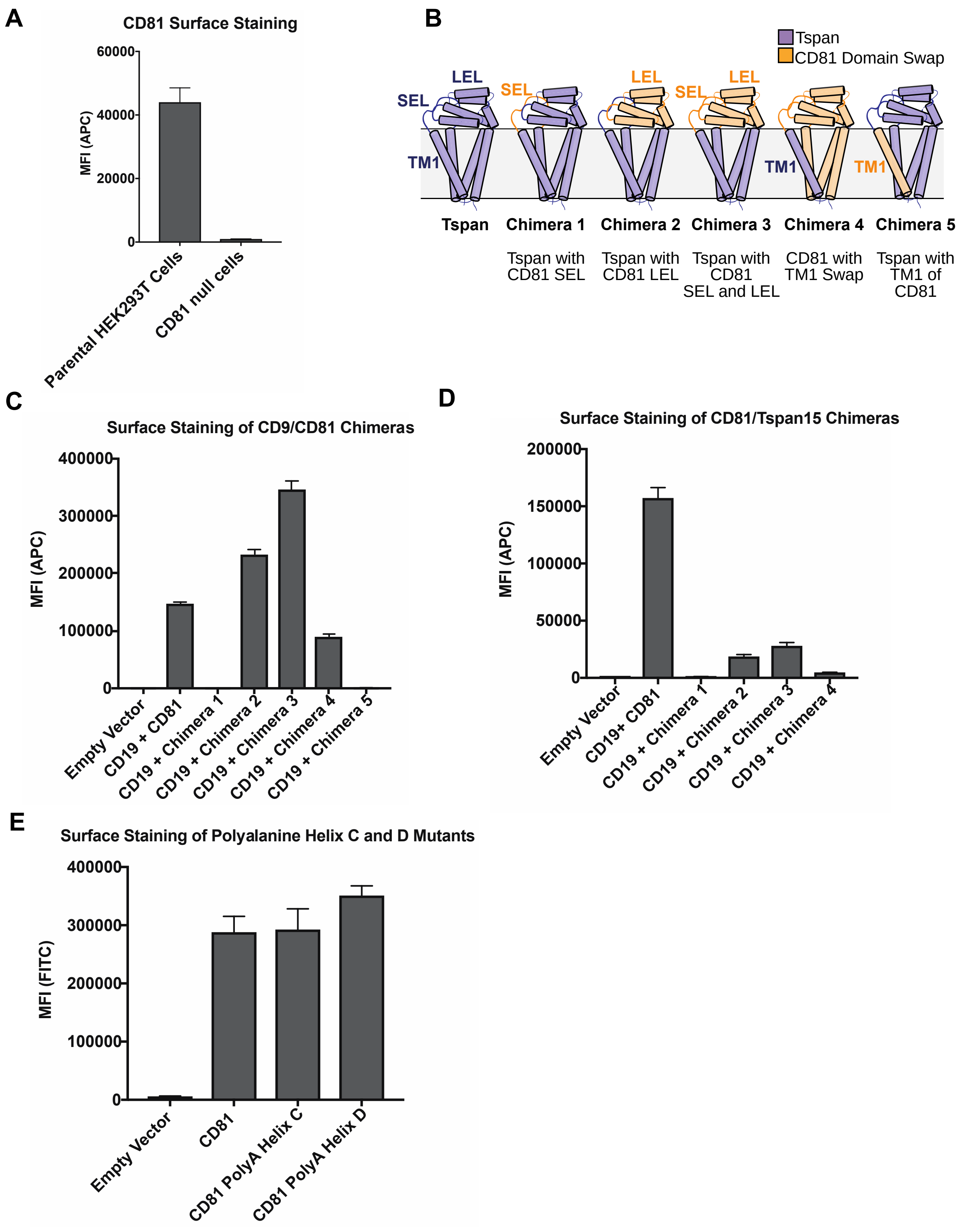


**Figure S1 (Related to Figure 1):** Surface staining of CD81 chimeras used in the CD19 Export Assay.

Expression was analyzed using an anti-CD81 antibody, so only chimeras with the large extracellular loop of

CD81 are detectable. **(A)** CD81 surface staining of parental HEK293T cells compared to CRISPR knockout cells. **(B)** Panel of CD81 chimeras used in export assay. (**C)** CD81 surface staining of CD9/CD81 chimeras detected with 5A6 antibody. **(D)** CD81 surface staining of CD81/Tspan15 C. elegans chimeras detected with 5A6 antibody. **(E)** CD81 surface staining of polyalanine mutants detected with Ab21. Error bars represent mean ± SEM of three independent experiments.

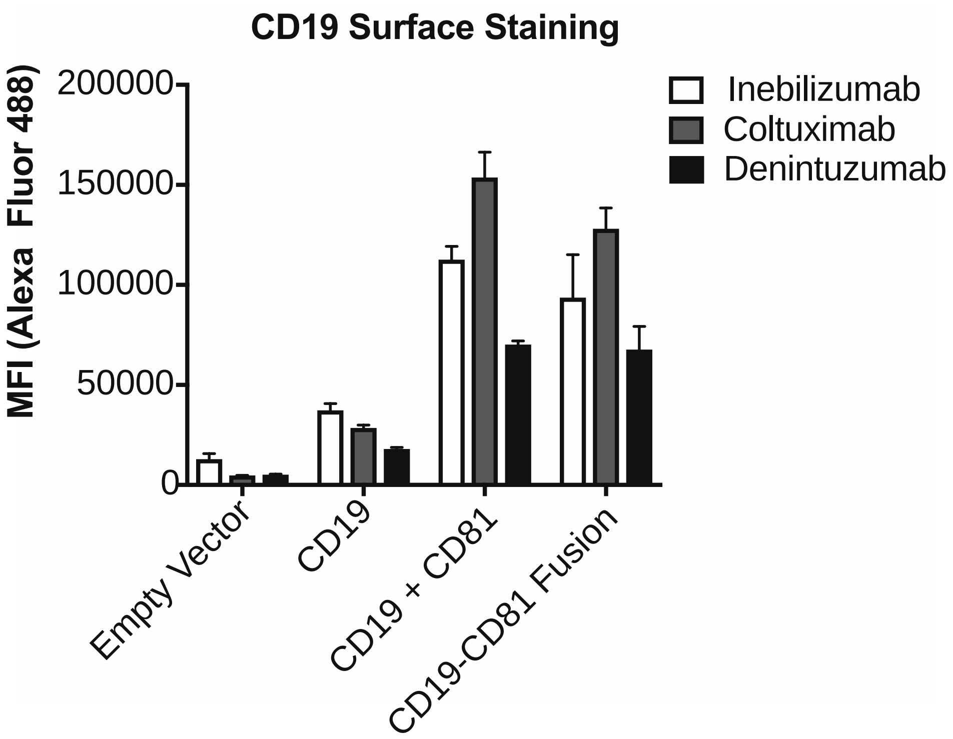


**Figure S2 (Related to Figure 2):** Validation of the CD19-CD81 fusion protein with a panel of CD19 antibodies. A panel of CD19 antibodies recognizes the CD19-CD81 fusion to the same degree as it recognizes wild type CD19, providing further evidence that the fusion protein is properly folded. Error bars represent mean ± SEM of three independent experiments.

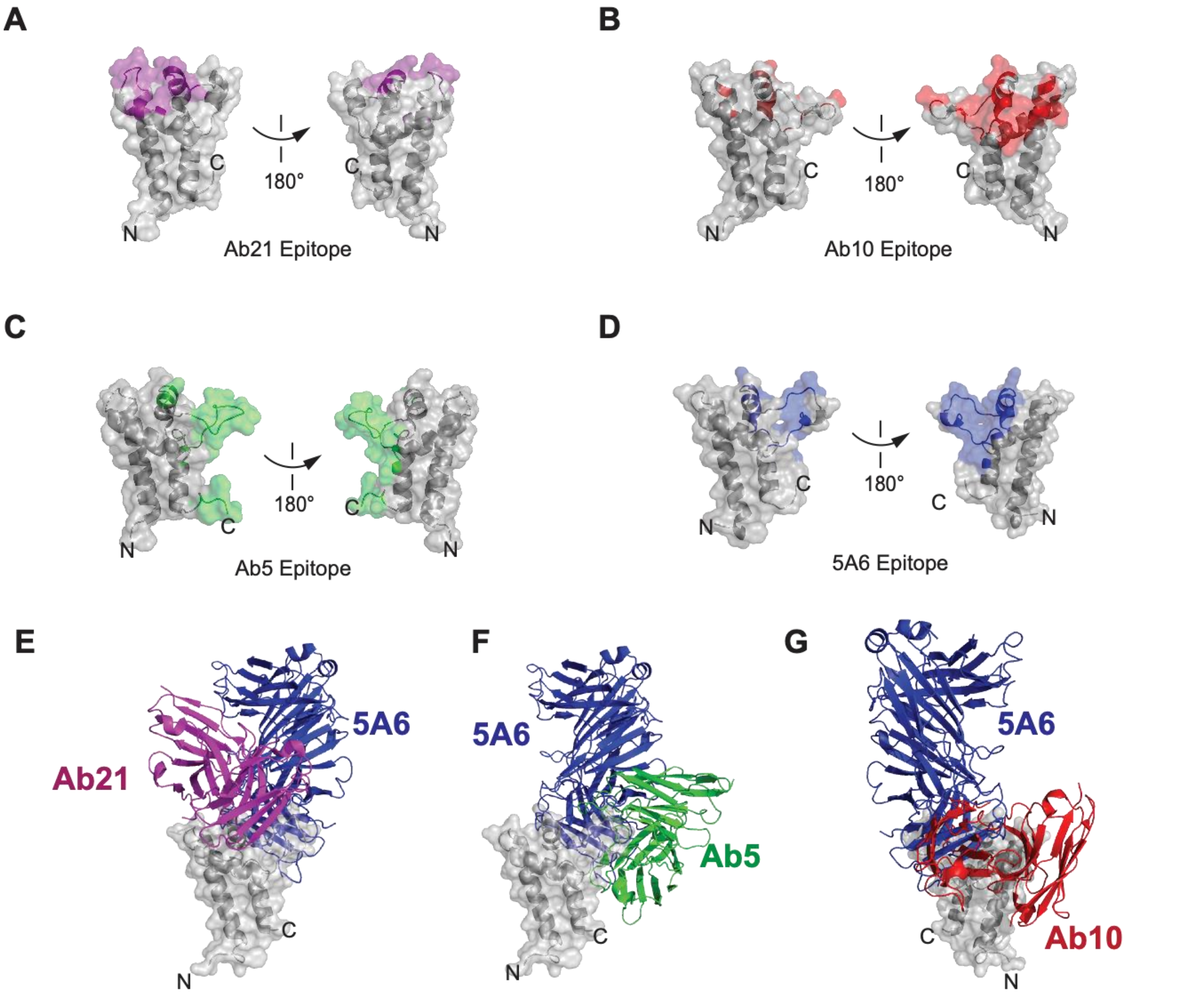


**Figure S3 (Related to Figures 2, 3, and 4):** Epitope comparison of Ab5, Ab10, Ab21, and 5A6. **(A)** Surface representation of CD81 large extracellular loop colored purple for residues within 4 Å of Ab21 (PDB 5DFW). **(B)** Surface representation of CD81 large extracellular loop colored red for residues within 4 Å of Ab10 (PDB 6EK2). **(C)** Surface representation of CD81 large extracellular loop colored green for residues within 4 Å of Ab5 (PDB 6EJM). **(D)** Surface representation of CD81 large extracellular loop colored blue for residues within 4 Å of 5A6. **(E)** Superposition of Ab21 and 5A6 complexes, comparing binding sites and angles of approach for Ab21 and 5A6. **(F)** Superposition of Ab5 and 5A6 complexes, comparing binding sites and angles of approach for Ab5 and 5A6. **(G)** Superposition of Ab10 and 5A6 complexes, comparing binding sites and angles of approach for Ab10 and 5A6.

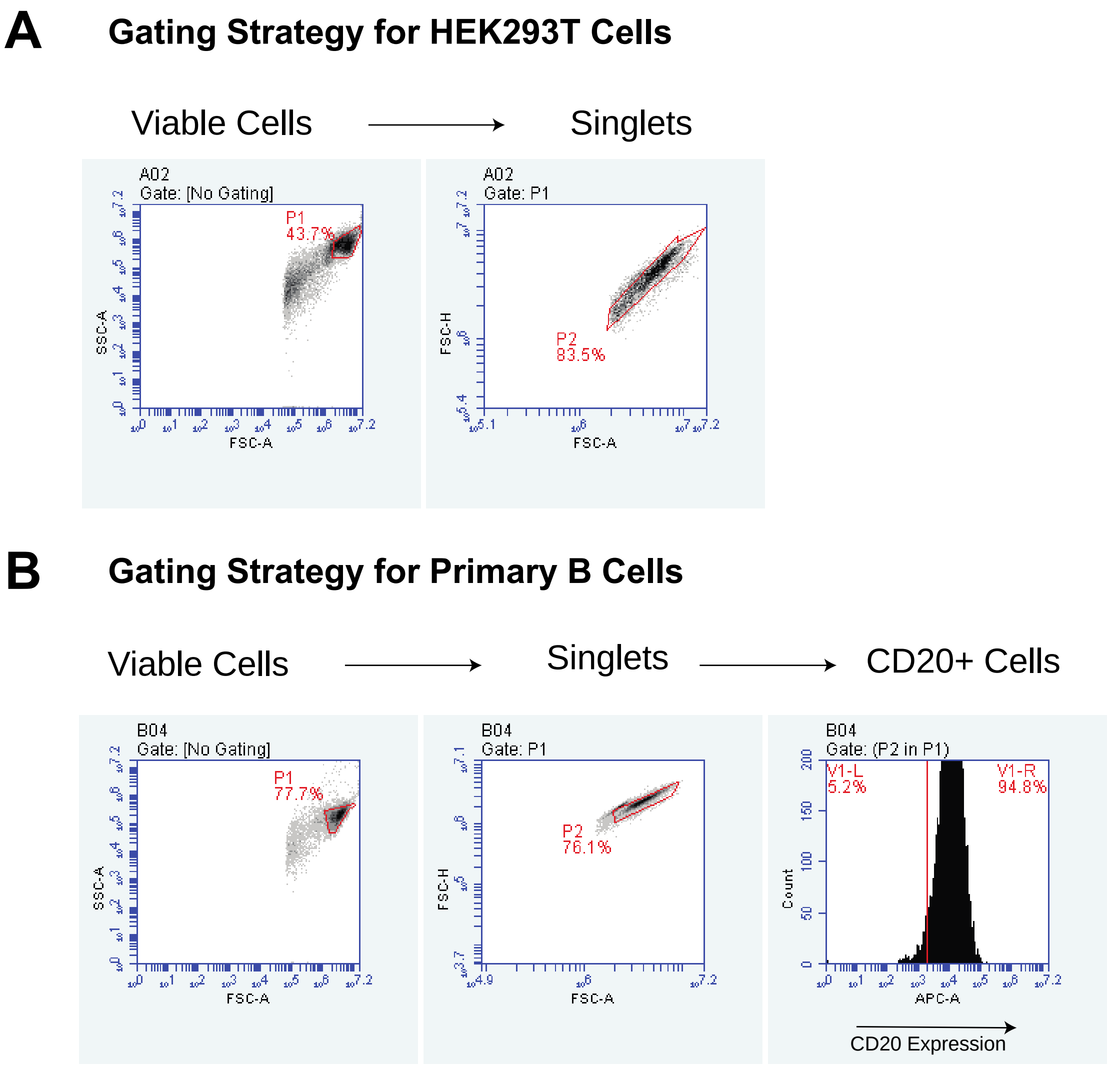


**Figure S4 (Related to Figure 1, Figure 2, and Figure 4):** Gating Strategy for CD81 null 293T cells and primary B cells **(A)** Representative gating strategy for CD81 null 293T cells used in the CD19 export assay and CD19-CD81 fusion protein validation experiments. Cells were first gated on live cells and then on singlet cells. **(B)** Representative gating strategy for primary human B cells. Cells were first gated on live cells, then on singlet cells, and then on CD20+ cells.

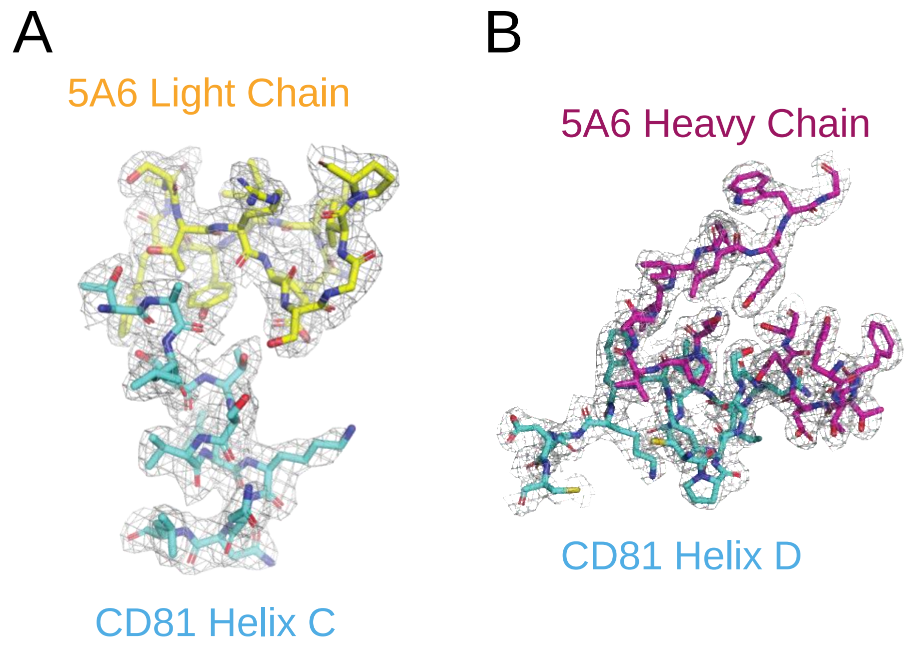


**Figure S5 (Related to Figure 3):** Representative Density in the CDRs of 5A6 $F_{ab}$. (A-B) Composite omit $2F_O$-$F_C$ electron density map contoured at 1.0 σ for CD81 Helix C (A) and CD81 Helix D (B).
